# Testing the evolutionary drivers of nitrogen-fixing symbioses in challenging soil environments

**DOI:** 10.1101/2022.09.27.509719

**Authors:** Carolina M. Siniscalchi, Heather R. Kates, Pamela S. Soltis, Douglas E. Soltis, Robert P. Guralnick, Ryan A. Folk

## Abstract

- While the importance of root nodular symbioses (RNS) in plants has long been recognized, the ecological and evolutionary factors maintaining RNS remain obscure. RNS is associated with environmental stressors such as aridity and nitrogen-poor soils; the ability to tolerate harsh environments may provide ecological opportunities for diversification, yet, nodulators are also diverse outside these environments.
- We test several environmental determinants of increased survival and enhanced diversification of RNS species, using an explicitly phylogenetic approach for the first time. We assembled the largest phylogeny of the nitrogen-fixing clade to date and a comprehensive set of abiotic niche estimates and nodulation data. We used comparative phylogenetic tools to test environmental and diversification associations.
- We found that RNS is associated with warm, arid, and nitrogen-poor habitats. However, RNS was gained long before lineages entered these habitats. RNS is associated with accelerated diversification, but diversification rates are heterogeneous among nodulators, and non-legume nodulators do not show elevated diversification.
- Our findings undermine the interpretation that RNS directly drove the invasion of challenging habitats, and do not support a direct relationship between soil or climate and the diversity of nodulators. Still, RNS may have been an important exaptation allowing further niche evolution.

## INTRODUCTION

Root nodule symbiosis (RNS), the encapsulation of endophytic nitrogen-fixing bacteria in specialized plant lateral root structures, is one of the most impactful forms of plant-microbe symbiosis in terrestrial landscapes, as measured by the scale of geographic terrain covered by nodulators (Tamme et al. 2021) and the physiological effectiveness of the process itself (Pankievicz et al. 2019). Intense research interest has focused on the phylogenetic origin (Soltis et al. 1995; Doyle 2011; Werner et al. 2014; Li et al. 2015; Kates et al. 2022) and functional genomic basis (Griesmann et al. 2018; van Velzen et al. 2018) of RNS itself, as well as understanding factors shaping current diversity and distributions (Sprent et al. 2017; Ardley and Sprent 2021). While the origin of RNS has recently attracted renewed interest (reviews: Sprent 2007; Doyle 2011; van Velzen et al. 2019), a link between the origins and modern prevalence of RNS remains poorly understood, leaving uncertain the ecological context that may have driven the gain of nodules, as well as how—or if—this gain enabled the ecological dominance of nodulators, and legumes in particular, in harsh landscapes.

All nodulators are restricted to the nitrogen-fixing clade (NFC), a group of four recognized taxonomic orders (Cucurbitales, Fabales, Fagales, and Rosales) including the legumes and nine other families (Soltis et al. 1995; The Angiosperm Phylogeny Group 2016). While species-level data are limited, based on our current understanding of the distribution of nodules (Afkhami et al. 2018; Tedersoo et al. 2018; Kates et al. 2022), approximately 52% of the species in the nitrogen-fixing clade are thought to have the ability to engage in RNS (Kates et al. 2022). The nodule organ is structurally diverse; nodulators have been classified based on their bacterial partners, either “rhizobia” (a broad term for bacterial partners of phyla Alphaproteobacteria and Betaproteobacteria, including the traditionally recognized genus *Rhizobium*, in which case the plant hosts are termed “rhizobial”) or *Frankia* (phylum Actinobacteria, with the plant hosts termed “actinorhizal”), a division that mostly follows host clade membership and morphological structure of the nodule itself (Pawlowski and Sprent 2008; Ardley and Sprent 2021).

Nodulators are both prevalent and diverse globally, and while many nodulating lineages are not especially species-rich, legumes are one of the largest and most prevalent plant families. This diversity has led to the interpretation of RNS as a potential “key innovation” (Koenen et al. 2013) that enabled the NFC to diversify broadly. However, spatial heterogeneity in the distribution of nodulators (Tamme et al. 2021) does not match the global distribution of nitrogen limitation (LeBauer and Treseder 2008), leading to questions about why nodulators are prevalent in landscapes that are not nitrogen-limited but absent from some nitrogen-limited habitats (Crews 1999). It remains unclear which environmental factors promote the presence of RNS (Pellegrini et al. 2016; Adams et al. 2016). A more complete understanding of the ecological pressures promoting RNS in present-day communities may provide insight into the original gain of the symbiosis, but these connections have not been thoroughly investigated because a framework for simultaneously incorporating evolutionary and spatial ecological knowledge is lacking.

Relatively few studies have attempted to understand the diversification and evolution of nodulating members of the nitrogen-fixing clade, with most focused on the legumes (Li et al. 2013; Koenen et al. 2013; Lavin et al. 2005; Sanderson and Wojciechowski 1996). Despite the popularity of “key innovation” arguments (Hunter 1998; Donoghue 2005; Li et al. 2013; Werner et al. 2014; Koenen et al. 2013), investigations of the nodule itself as a driver of diversification have rarely been attempted (Afkhami et al. 2018). This paucity of work relates partly to the challenge of gathering information about an underground trait across large numbers of species and a wide geographical distribution (reviewed in Sprent 2009), a challenge at least partially addressed by several databasing efforts (Werner et al. 2014; Afkhami et al. 2018; Tedersoo et al. 2018; Tamme et al. 2021; Kates et al. 2022).

The largest-scale attempt to date on understanding the relationship between environmental factors and the diversification of nodulators (Tamme et al. 2021) aimed at identifying drivers of species richness in clades of plants with RNS. Tamme et al.’s (2021) primary finding was that bacterial partner choice influenced different relationships with latitude, with rhizobial taxa associated with the periphery of the tropics and actinorhizal taxa associated with high-latitude and montane environments. Aside from latitude and elevation, climate and soil were evaluated as drivers of these geographical disparities jointly using an ordination approach.

Although Tamme et al. (2021) invoked evolutionary theory, that study lacked a phylogenetic framework that could address this hypothesis. Here we instead directly incorporate evolutionary information, such as the timing of the origin(s) of the symbiosis itself, and the long-term history of how lineages switched to different habitats to make a stronger evolutionary argument linking nodules to the environments in which they are often found. We focus specifically on estimating niche occupancy and diversification rates, and tested two basic and closely connected hypotheses about the ecological attributes associated with the macroevolutionary success of the nitrogen-fixing clade, as quantified in terms of lineage diversity and niche occupancy. The first hypothesis is that nodulating species are more prevalent than non-nodulating species in nitrogen-poor and arid niches, and the second is that RNS is associated with shifts into arid and nitrogen-poor habitats and accompanying ecological release (i.e., increased rates of habitat evolution). As corollaries, we tested for three additional expected outcomes of ecological release due to RNS: (3) in the nitrogen fixing clade, nodulating species have higher diversification rates than non-nodulating species; (4) correspondingly, diversification rate regimes are directly connected with RNS and its environmental drivers, viz., with the entry of nodulators into differing, more challenging limited nitrogen and high aridity niches, and, (5) nodulators have greater niche breadth reflective of ecological release, as well as niche occupancy of areas with greater nitrogen limitation than their close relatives.

## METHODS

### Circumscription

Our focus was on the nitrogen-fixing clade as defined by Soltis et al. (1995), excluding angiosperms with other biologically distinct symbiotic organs and forms of the symbiosis that do not involve specialized organs, or “associative” nitrogen fixation (*sensu* Pankievicz et al. 2019). This focus follows other articles that show different spatial and environmental patterns of evolutionarily and ecologically distinct symbiotic plants outside the nitrogen-fixing clade (Tamme et al. 2021), as well as important differences in physiological efficiency (Pankievicz et al. 2019) and a probable lack of homology (Doyle 2011, 2016).

### Supermatrix assembly and processing

To date, large-scale investigations of nodulators have either relied on DNA supermatrices that generated trees with relatively poor support (Werner et al. 2014) or did not include a phylogenetic framework (Tamme et al. 2021). Here, we improved phylogenetic estimates of the nitrogen-fixing clade through an integration of available GenBank data with a recently reported major phylogenomic analysis (Kates et al. 2022) to maximize species presence in the ultimate phylogenetic product.

First, we assembled a matrix from available GenBank data using PHLAWD (Smith et al. 2011). Following a trial of clustering and reference-based methods (see Smith and Walker (2019), we selected reference-based methods, because they resulted in high per-locus taxon coverage. We handled high intralocus divergence using profile alignment methods as detailed below. Based on taxon occupancy, the following loci were collected from the nuclear ribosomal RNA cistron: 18S rDNA, 28S rDNA, and the external and internal transcribed spacers (ETS and ITS, respectively); the low-copy portion of the nuclear genome: *gapc, phyc*; the chloroplast genome: *atpB-rbcL, atpB, matK, ndhF, psba, rbcL, rpl20-rps12, rpoC1, rps16, trnD-trnY-trnE, trnL-F, trns-trng*; and the mitochondrial genome: *matR, nad1*.

To improve taxon representation across loci, we then assembled the same loci from off-target portions of capture data from the (Kates et al. 2022) dataset using aTRAM2 (Allen et al. 2018) and a representative sequence for each locus from each taxonomic order as references. Five assembly iterations were used to optimize assemblies, and preliminary orthology was assessed based on bitscore with the top hit taken. These assemblies were merged with PHLAWD results for alignment.

Due to the use of variable loci across a wide scope of phylogenetic comparisons, particularly ITS (McMahon and Sanderson 2006; Soltis et al. 2013; Sun et al. 2016, 2019), we used a profile alignment strategy to improve alignments. Each locus was aligned in MAFFT within data subsets by clades recognized at the family rank (The Angiosperm Phylogeny Group 2016). Alignments were then merged across families with the MAFFT --merge function. Thus, alignments were ultimately reunified to render a single locus from the point of view of the final analysis. The final concatenated alignment was 51,868 bp in length for 16 loci with 84.9 percent missing data and 19,756 species.

For the tree search, we used RAxML-NG (Kozlov et al. 2019). The matrix was treated as a single partition, and the character evolution model was chosen with the --parse function in RAxML-NG. We imposed a constraint backbone containing fully resolved clades based on the tree of (Kates et al. 2022), so that the backbone was inferred by phylogenomic data, and GenBank data were used primarily for extending taxon sampling. The tree search was implemented for individual orders or clades (i.e., Fabales were split into five clades and Rosales were split into two, given their richness) and stitched to a backbone inferred from 100 representative taxa.

We manually reviewed and excluded taxa with excessively long branches and tips that violated family monophyly (subfamilial monophyly in larger families), using higher-level taxonomies in the World Checklist of Vascular Plants (WCVP; https://wcvp.science.kew.org/) and family-specific resources for Fabales (Lewis 2005, LPWG 2017). Remaining taxon duplicates after synonymization were resolved by expected phylogenetic position based on previous studies or randomly in the case of ties. The final tree product has 54.6% species-level taxon coverage (16,375 species out of ∼30,000; Kates et al., 2022). Compared with previous phylogenetic investigations of the nitrogen-fixing clade, e.g., seven loci with 13.0% species-level coverage (Zanne et al. 2014; Werner et al. 2014; Menge and Crews 2016) and three loci with 3.1% coverage (Li et al. 2015), this is the most comprehensive phylogeny to date in terms of species richness and a 28% increase (3,607 species) in sampling compared to (Kates et al. 2022), which sampled a partly overlapping set of 12,768 species for 86 loci.

Divergence time estimation used penalized likelihood as implemented in TreePL (Smith and O’Meara 2012). Eleven secondary calibration points were used following (Magallón et al. 2015), primarily guided by limitations in taxon sampling and differing topology in a few cases. The “prime” option was used to calculate the optimal parameters for the analysis, which was conducted with random cross validation.

### Nodulation data

Previous efforts to assess RNS at large scales have primarily relied on collation of reports of nodule presence, including variable levels of empirical evidence (Werner et al. 2014; Tedersoo et al. 2018). Here, we use a recent comprehensive nodulation database effort that completely reevaluated the literature on nodulation states in the nitrogen-fixing clade using high standards of evidence (Kates et al. 2022).

### Environmental data acquisition

Occurrence records were downloaded from both iDigBio and GBIF (R packages rgbif and ridigbio). We queried all species sampled in GenBank and Kates et al. (2022) with an occurrence limit of 1,000 and hasCoordinate = TRUE. We increased species representation of records by taking advantage of the fact that collectors often report numerous species from single collecting sites, but often only a subset of co-located organisms are georeferenced. Using a complete download of plants on GBIF, we utilized a prototype gazeteer of descriptive localities and associated georeferences (Zermoglio et al. in review; https://github.com/VertNet/bels). We then applied this gazeteer, using its best practices-based matching algorithm, and populated latitude and longitude fields in cases where textual localities matches were found. We implemented several steps of occurrence record curation with the overall goal of retaining records conformant to expert-determined species ranges and without other clear georeferencing issues. These steps were executed on a per-species basis as follows: first, we removed all exact duplicates and all GPS records only determined to the nearest integer.

We then removed all outliers beyond three standard deviations from the Euclidean centroid of the occurrence records. Using the World Checklist of Vascular Plants via the POWO API (Python library pykew; https://pypi.org/project/pykew/), we matched taxa to occurrence data and removed all occurrence records outside of the level-three TDWG regions considered as native occurrences in POWO. Finally, for environmental data extractions, per-pixel duplicates were removed to reduce biased oversampling of sites due to differences in collecting effort or spurious overcounting due to specimen duplicates. In total, 24,940 species were retained after filtering with georeferenced records, and 22,711 remained after excluding species with unknown/variable scorings for the nodulation trait. Inclusion of gazeteer data resulted in an average increase of 154 records per species and an average record growth of 39.8%.

We used environmental data assembled from WorldClim v1 (Hijmans et al. 2005), *n* = 19) and ENVIREM (from which the UNEP aridity index was calculated; (Title and Bemmels 2018), SoilGrids250m (Hengl et al. 2017), land-cover data (Tuanmu and Jetz 2014), and a DEM (Digital Elevation Model, from which were derived slope and aspect; GTOPO30; https://www.usgs.gov/centers/eros/science/usgs-eros-archive-digital-elevation-global-30-arc-second-elevation-gtopo30). From these environmental data, we selected 16 variables that (1) were not highly collinear with one another across the global study area (Pearson’s *R*^2^ < 0.7) and (2) were biologically relevant considering our research questions. The climatic variables included were mean annual temperature (BIO1), isothermality (BIO3), temperature seasonality (BIO4), mean annual precipitation (BIO12), precipitation seasonality (BIO15), precipitation of driest quarter (BIO17), and aridity. Topographic and soil variables used were elevation, soil nitrogen content, soil carbon content, soil pH, soil coarse fragment content, and soil sand content. To capture overall vegetative context, we included land-cover by deciduous broadleaf, herbaceous, and coniferous (“needle-leaf”) vegetation. For multispecies analyses, we summarized the niche occupancy of each species by its per-variable means and niche breadth (see below for niche breadth calculation).

### Name resolution and taxon matching

Names from each data source were reconciled against WCVP to update synonymies and generic re-circumscriptions. After name reconciliation, 13,244 species were successfully matched between phylogenetic and environmental data.

### Niche breadth

Niche breadth captures overall ecological tolerances of species and can be interpreted as a measure of ecological specialization (Sexton et al. 2017). It has been suggested both from the perspective of the bacterial symbiont (Masuda et al. 2020; Criminger et al. 2007) and plant host (Xu et al. 2021) that nitrogen-fixing strategies increase niche breadth, a hypothesis yet to be tested at this broad scale. Niche breadth was calculated using hypervolume concepts (Hutchinson 1957) by calculating per-species ranges for each environmental variable, normalizing to mean 0 and standard deviation 1, and multiplying across variables. We used this niche breadth measure as the response variable in a set of linear models to ask whether: (1) nodulators have higher niche breadth, consistent with an association with weedy/generalist strategies, or (2) independent of nodulation, higher niche breadth is associated with low nitrogen or high aridity.

### Model fitting

We first fit logistic regression models to environmental predictors, treating nodulation state (present versus absent) as a factor to ask whether nodulators differentially occupy niche space compared to non-nodulators. We then used a separate model set based on phylogenetic generalized least squares (PGLS) to control for non-independence of environmental variables due to shared evolutionary history, considering both models with and without evolutionary relationships to understand how phylogenetic structure may have influenced previous results.

Uncorrected logistic regression and PGLS approaches represent alternative philosophies with differing interpretations relevant to the field of inquiry (Guillerme et al. 2020; Westoby et al. 1995). For instance, asking whether gain of nodulation was associated with a shift into arid habitats requires phylogenetic correction (Felsenstein 1985) because phylogenetic non-independence is clearly relevant to a question framed in evolutionary terms. By contrast, asking the simpler question of whether there are more nodulators in deserts than in other regions does not require phylogenetic correction (de Bello et al. 2015) because learning that all desert nodulators are closely related would be interesting, and potentially tied to different historical and current processes, but irrelevant. As our analysis represents the first time PGLS has been applied to assess the environmental niche of nodulators, both analyses are needed to fully contextualize any differences in interpretation from previous results. The R package caper (Orme et al., 2018) was used to assemble a variance-covariance matrix associating all tree tips with their corresponding environmental variable values, calculated as explained above. We then fit both individual models for each variable and one combined model containing all environmental variables.

In addition to the environmental models above, we fit a series of evolutionary models of environmental characteristics to obtain estimates of evolutionary rates and root states using the R package geiger (Harmon et al. 2008; Pennell et al. 2014). A lower rate of niche evolution in nodulating vs. non-nodulators would suggest that the environments preferred by nodulators represent an evolutionarily stable strategy; conversely, a higher rate of niche evolution implies that nodulators shift in niche frequently and are not specialized in any ancestral habitat. We used standard model comparison techniques with AIC_c_ for model choice, fitting Brownian, Ornstein-Uhlenbeck (OU), and Early-Burst models. In each case, to gain insight into nodulation-dependent environmental interactions, we segregated the tree and environmental data into nodulating and non-nodulating portions and fit separate models, performing model selection separately by data partition.

Since we were interested in the biological interpretation of rate estimates from the foregoing models, we implemented a second model comparison framework to test whether nodulators and non-nodulators differ in evolutionary rate. To do this, we estimated a constrained OU model for nodulating taxa, parameterizing evolutionary rate as identical to that estimated for non-nodulators. We compared this to the unconstrained model with a likelihood ratio test to determine if rates were nonequivalent between nodulators and non-nodulators. Finally, we performed assessments of niche conservatism in the environmental variables using the lambda test in the R package phytools (Revell 2012). Multiple comparisons were corrected by the Hochberg method (Hochberg 1988).

### Assessment of speciation rates

Evolutionary gain of nodules may be associated with downstream evolutionary radiations, given the potential for a selective advantage of nodulators in challenging environments. Examination of this question has so far proved inconclusive (Afkhami et al. 2018), but previous efforts used methods that are potentially problematic (Rabosky 2018) with highly incomplete phylogenies. We used measures of speciation assessed at the present (so-called “tip-rates”) for their straightforward interpretation, relative robustness to speciation-extinction model violations, and better agreement with alternative diversification measures from the fossil record (Title and Rabosky 2019; Louca and Pennell 2020; Upham et al. 2021). DR rates were calculated on the timetree above using previously published R code (Sun et al. 2020), and a test of the relationship between nodulation and speciation rates was implemented in FiSSE (Rabosky and Goldberg 2017) to account for evolutionary pseudoreplication.

## RESULTS

### Hypothesis 1, root nodule symbiosis and environmental context

We first asked whether nodulating and non-nodulators differ in environmental occupancy using generalized linear models and found that nodulators are in soils with significantly lower nitrogen content than non-nodulators in the nitrogen-fixing clade (*p* < 2e-16). Based on variance explained (univariate adjusted *R*^2^ > 0.05; Table 1), the five most important factors in determining environmental occupancy of nodulators were (in descending order) aridity, carbon, nitrogen, precipitation (BIO12), and precipitation of driest quarter (BIO17). Hereafter, we will focus on seven variables, including these five as well as mean annual temperature (BIO1) and temperature seasonality (BIO4). Among these seven, aridity had the greatest explanatory power (assessed based on normalized coefficients; adjusted *R*^2^ was also highest at 0.1109); the overall explanatory power of a combined MANOVA model was moderate (*η*^*2*^ = 0.1637). However, segregating these results by RNS type (rhizobial vs. actinorhizal) shows phylogenetic heterogeneity. Rhizobial taxa are most distinct in their soil and precipitation niche. Actinorhizal taxa differed from rhizobial and non-nodulators primarily by aridity, temperature, and elevation niche (Fig. 1). Rhizobial nodulators drive the difference in nitrogen niche observed for nodulators overall, while actinorhizal taxa and non-nodulators have near-identical nitrogen niche. Consistent with heterogeneity between rhizobial and actinorhizal taxa, actinorhizal and rhizobial RNS taxa show differences at lower phylogenetic levels (probability densities plotted in Fig. 1).

**Table 1.** Environmental model results for linear and PGLS models. Highlighted cells were those with *R*^2^ exceeding 0.05 and two further additional found to be important in previous studies, which were subject of closer focus (see main text).

| Dependent variable: | Standardized coefficient | $R^2$ (linear) | $p$-value (linear) | $R^2$ (PGLS) | $p$-value (PGLS) |
| --- | --- | --- | --- | --- | --- |
| aridity | -0.358 | 0.116 | < 2.2e-16 | -6.00E-05 | 0.7109 |
| bio15 | 5.932 | 0.014 | < 2.2e-16 | -6.95E-05 | 0.9825 |
| bio17 | -56.161 | 0.067 | < 2.2e-16 | -6.74E-05 | 0.8633 |
| bio3 | -0.092 | -0.0001 | 0.0001682 | -6.94E-05 | 0.9761 |
| bio4 | -37.592 | -0.00003 | 0.05853 | -5.70E-05 | 0.672 |
| carbon | -14.161 | 0.098 | < 2.2e-16 | -6.95E-05 | 0.9975 |
| coarsefragment | -0.228 | 0.0003 | 0.9809 | -6.11E-05 | 0.7276 |
| deciduousbroadleaf | -0.063 | -0.00005 | 0.2141 | -6.60E-05 | 0.8217 |
| elevation | -56.716 | 0.002 | 0.03519 | -6.78E-05 | 0.8751 |
| herbaceous | 1.734 | 0.008 | < 2.2e-16 | -5.97E-05 | 0.7066 |
| needleleaf | -0.999 | 0.004 | 6.378e-08 | -5.86E-05 | 0.6926 |
| nitrogen | -53.116 | 0.084 | < 2.2e-16 | -3.11E-05 | 0.4574 |
| ph | 3.297 | 0.022 | < 2.2e-16 | -4.50E-05 | 0.5525 |
| precipitation | -375.420 | 0.07 | < 2.2e-16 | -6.04E-05 | 0.7179 |
| sand | 4.228 | 0.027 | < 2.2e-16 | -5.17E-05 | 0.6126 |
| temperature | 4.758 | 0.001 | 0.3559 | -5.47E-05 | 0.6441 |

**Fig. 1.**
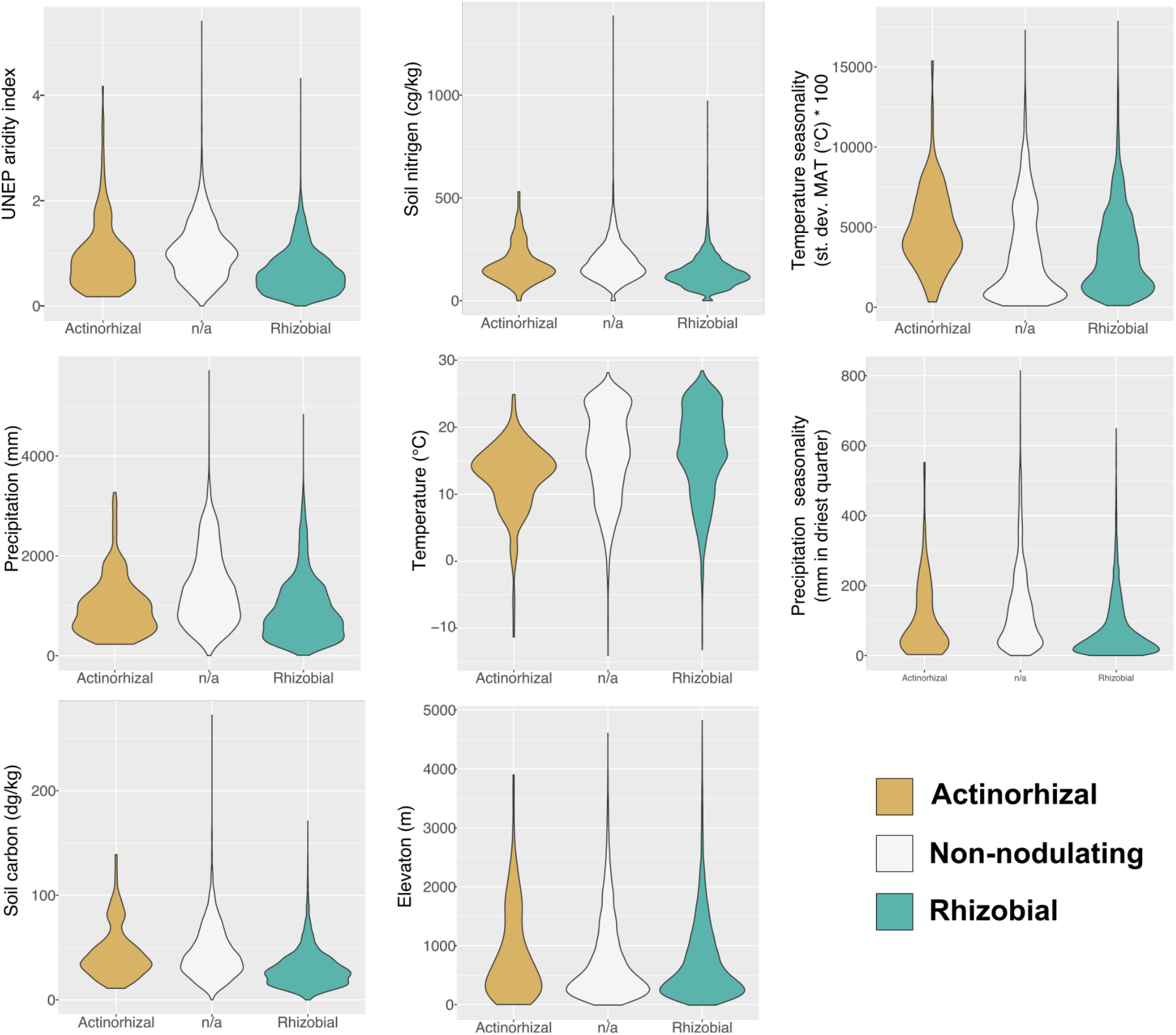
Violin plots of environmental attributes against nodulation status and type, focusing on the seven main predictors as identified in the main text, as well as elevation as identified in previous literature. Rhizobial taxa are most distinct in their soil and precipitation niche; actinorhizal taxa are markedly different in aridity, temperature, and elevation. Note the complex multimodality nodulators often show, due to differences among subclades, and that actinorhizal and rhizobial forms of symbiosis do not necessarily show similar environmental responses.

When we used PGLS approaches to account for phylogenetic signal, we found that significance disappeared for all environment-RNS correlations, with near-zero *R*^2^ values (Table 1). Therefore, phylogenetic co-structuring of environmental preferences and RNS is responsible for niche differences of nodulators, limiting our ability to invoke at least a direct causative role.

### Hypothesis 2, ecological release and habitat shift

Due to the importance of phylogenetic signal shown in the PGLS results, we further investigated the evidence for niche conservatism in RNS using the lambda test, focusing on the seven main environmental predictor variables. All tested factors were significant and similar in estimates of the magnitude of signal (Table 2), suggesting the importance of niche conservatism in the nitrogen-fixing clade.

**Table 2.**
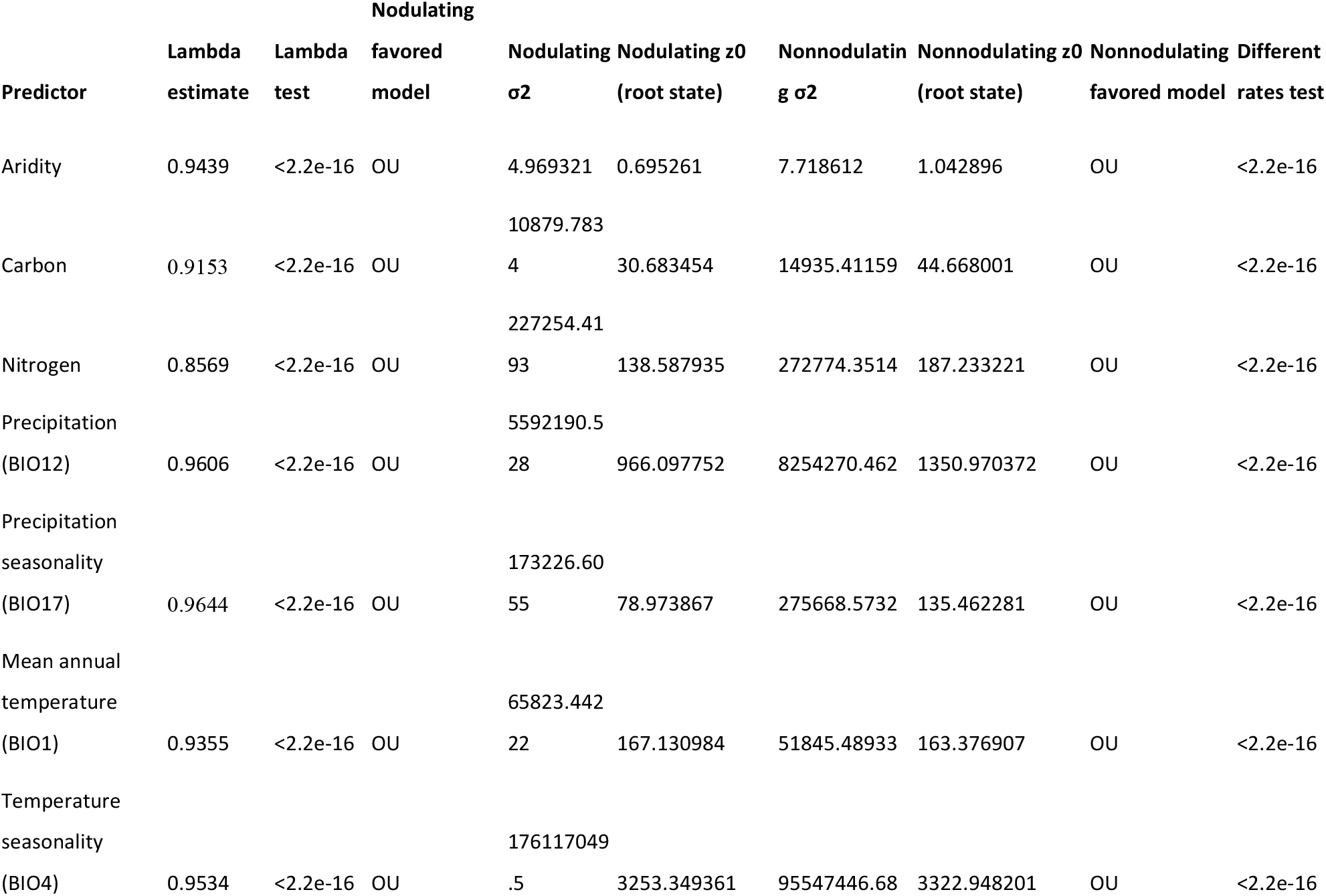
Phylogenetic signal and evolutionary model results. λ = 1 corresponds to Brownian expectations of phylogenetic structure and λ = 0 corresponds to a complete lack of phylogenetic structure to niche occupancy; see (Wiens and Graham 2005; Cooper et al. 2010; Wiens 2008; Losos 2008)). *σ*^2^ represents evolutionary rate scaled to trait units; *z*_0_ represents the estimated root state. Here *p-*values are corrected by the Hochberg method.

We then fit a series of evolutionary models to ask whether nodulating and non-nodulators differ in niche evolution rates, investigating the same seven predictors as above. We subset the tree to its nodulating and non-nodulating portions and used the models to examine estimated root states (interpretable as a phylogenetically weighted average of tip states) and evolutionary rates. Based on model tests, OU models were favored for all comparisons (Table 2). For five of the variables (aridity, nitrogen, carbon, precipitation [BIO12], and precipitation seasonality [BIO17]), we found significantly *lower* rates of niche evolution in nodulators, corresponding to the concept of RNS as a stable evolutionary strategy. Specifically, nodulators occur in, and infrequently leave arid ancestral environments with less fertile soils and lower precipitation than non-nodulators. Consistent with this result, mapping niche space vs. nodulation states (Fig. 3) revealed nested habitat transitions within clades fixed for nodulation state. However, temperature and temperature seasonality, which were not among the most important variables for contemporary species, are associated with higher evolutionary rates in nodulators, with somewhat higher ancestral temperature and lower ancestral temperature seasonality than non-nodulators. Rate differences suggest these temperature attributes represent an unstable ancestral niche, consistent with results (above) for extant species. Based on a model comparison framework, the rate differences we recovered were highly significant (*p* < 2e-16 in all cases after correction for multiple comparisons).

### Hypothesis 3, speciation rates and root nodule symbiosis

We found that nodulators indeed have significantly higher speciation rates than non-nodulators (FiSSE *p* = 0.042; speciation ∼3-fold higher for nodulators; Fig. 2a); since FiSSE accounts for evolutionary pseudoreplication (Rabosky and Goldberg 2017), this relationship could be taken as potentially causal. However, distinguishing between actinorhizal and rhizobial RNS reveals that this diversification effect is driven by only a few key clades: rhizobial nodulators have a long right-hand tail that includes essentially all species in the nitrogen-fixing clade with the highest rates of speciation (Fig. 2b). By contrast, actinorhizal plants have mean speciation rates slightly *lower* than non-nodulators (Fig. 2b), suggesting different diversification dynamics that might relate to the nature of the symbiosis with partner bacteria. Likewise, diversification is non-uniform within the legumes, with the highest rates in caesalpinioid legumes (FiSSE *p* = 0.03; speciation ∼3-fold higher for nodulators within this clade than for non-nodulators; Fig. 2c), while papilionoid legumes show fewer rapid diversifiers despite their greater species richness.

**Fig. 2.**
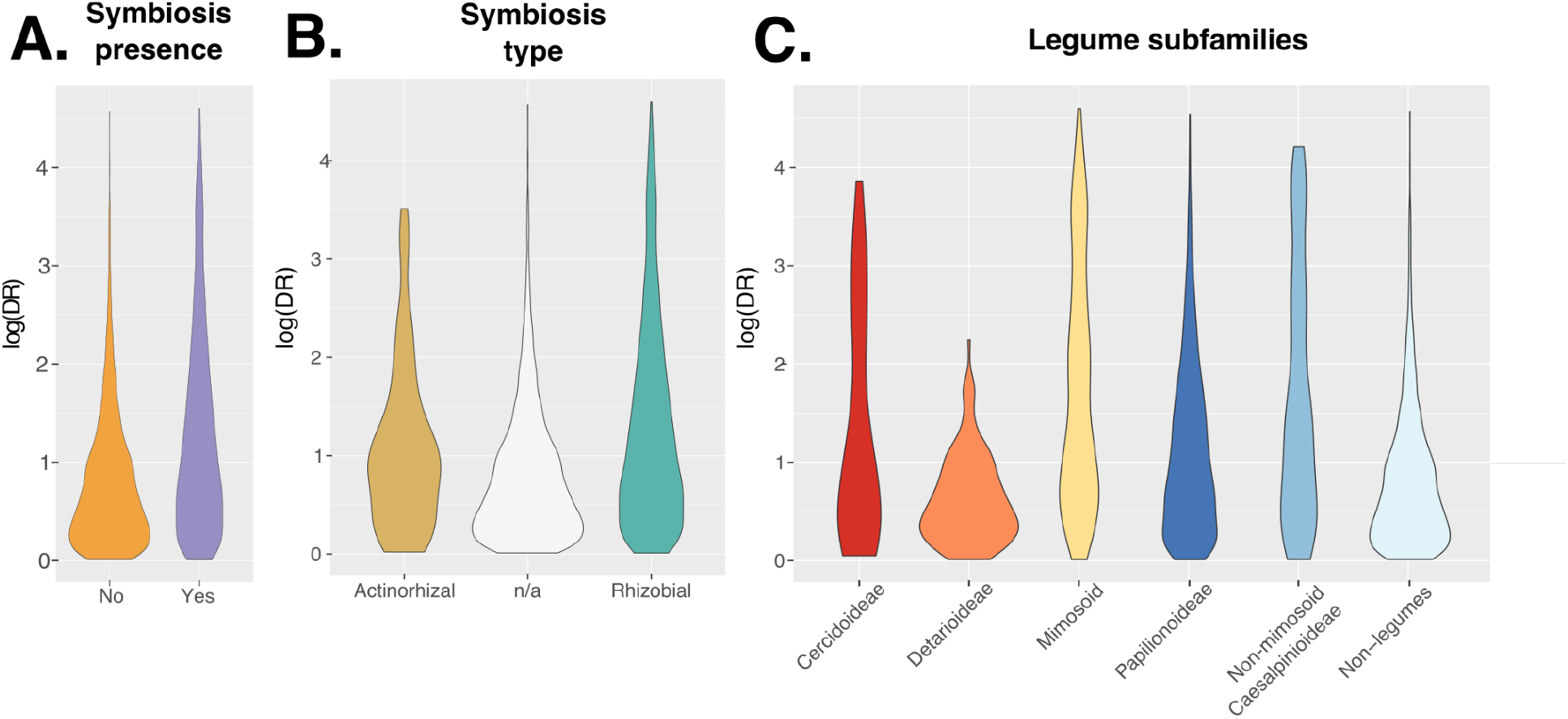
Violin plots of speciation (DR) against nodulator state. (A) DR vs. presence-absence of nodules. (B) DR vs. presence-absence of nodules, with presence subdivided into actinorhizal and rhizobial subgroups. (C) DR vs. legume subfamily, showing high rhizobial diversification rates are not unique to nodulating legumes and occur mostly in subfamily Caesalpinioideae, with the highest rates seen in the mimosoid clade of the subfamily.

### Hypothesis 4, speciation rates and habitat

We investigated whether disparities in diversification rate were associated specifically with differences in precipitation and soil regimes. To investigate the most challenging environments among the predictors included here, we segregated environments using UNEP definitions of arid (aridity index ≤ 0.05) and non-arid environments (aridity index > 0.05) and segregated soil nitrogen conditions by sites below the lower 5th percentile (low nitrogen) and conditions above this threshold (high nitrogen). Using FiSSE again, we tested whether these extreme climatic and soil conditions were associated with speciation rate. This comparison was non-significant (aridity: *p* = 0.2987; nitrogen: *p* = 0.4845); likewise, dividing this comparison into rhizobial and actinorhizal RNS also showed non-significance (all *p* > 0.2). Hence, some nodulators show increased speciation rates relative to non-nodulators within the nitrogen-fixing clade, and nodulators disproportionately occupy marginal areas. However, elevated diversification rates are not specific to the most marginal habitats that nodulators occupy, and therefore rapid diversification cannot be attributed to ecological release into these most challenging environments.

### Hypothesis 5, increased nodulator niche breadth

We found that nodulators do not differ from non-nodulators in niche breadth (linear model, *p =* 0.4958), also inconsistent with the ecological release hypothesis. Likewise, aridity and nitrogen niche are not associated with niche breadth (both *p* > 0.1, non-native range excluded), suggesting a general lack of any differences in niche breadth association with nodulating ecologies.

## DISCUSSION

### Root nodule symbiosis and harsh habitats

RNS, a trait linked to physiological performance advantages in challenging environments, should have a profound effect on the host plant niche. Numerous studies have focused on how environmental niche associates with RNS, whether in legumes (Pellegrini et al. 2016; Doby et al. 2022), broader groups of unrelated symbionts (Tamme et al. 2021), or various subclades, functional classes (particularly trees: (Steidinger et al. 2019; Menge et al. 2019), or habitats (Jin et al. 2013). Any climatic relationship found with RNS will beg the second question: why does RNS favor survival or preponderance in particular environments? Such hypotheses have been divided between those that alternatively view nodulator niche in terms of either present-day environment and those that attempt to link to historical constraints rooted in the origin of RNS (reviewed in Crews 1999). A general lack of direct phylogenetic approaches in most previous investigations (but see Menge and Crews 2016) has hindered making broader conclusions or integrating across these potential explanations.

Our primary focus was on testing evolutionary explanations more closely in the context of data and methods commonly employed to test ecological explanations. Our first and most basic result was that our analyses that do not remove phylogenetic autocorrelation corroborate recent results (Doby et al. 2022), where aridity is the most important predictor of nodulator environment occupancy, followed by soil and precipitation attributes. However, after accounting for phylogenetic signal, the significance of the relationship effectively disappears, underlining the importance of considering evolutionary pseudoreplication before attempting evolutionary extrapolation (Tamme et al. 2021).

While both RNS and environmental niche are highly conserved in the nitrogen-fixing clade, we find a differing evolutionary timeline for the gain of nodulation and the shift into environments that favor them, suggesting an indirect relationship between the two. Mapping environment and nodulation states (Fig. 3) shows that clades containing RNS shifted into favorable environments long after the gain of RNS itself. The earliest history of RNS has proven controversial (Doyle 2011; Werner et al. 2014; van Velzen et al. 2018; van Velzen et al. 2019; Kates et al. 2022), but it likely originated early in the evolutionary history of the nitrogen-fixing clade (Sprent 2007). Shifts into arid environments occurred far later in the history of the clade, consistently with current understanding of the timeline of aridification in today’s landscapes (Pound et al. 2012; Zachos et al. 2001; Potter and Szatmari 2009; Li et al. 2013; Sprent et al. 2017). In summary, while nodulators differ from non-nodulators within the nitrogen-fixing clade in their specialization to marginal habitats, the origins of RNS far predate shifts into these habitats and therefore cannot be invoked as directly causative, as previously argued by (Sprent 1994a). However, the fact remains that contemporary nodulators and non-nodulators in the nitrogen-fixing clade show stark differences in the environments they occupy today. In contrast to the traditional “key innovation” explanation (Koenen et al. 2013; Doyle 2011; Doyle and Luckow 2003), interpreting nodulation as an ecological exaptation (Bock 1959; Gould and Vrba 1982; Jacob 1977) is consistent with these results.

**Fig. 3.**
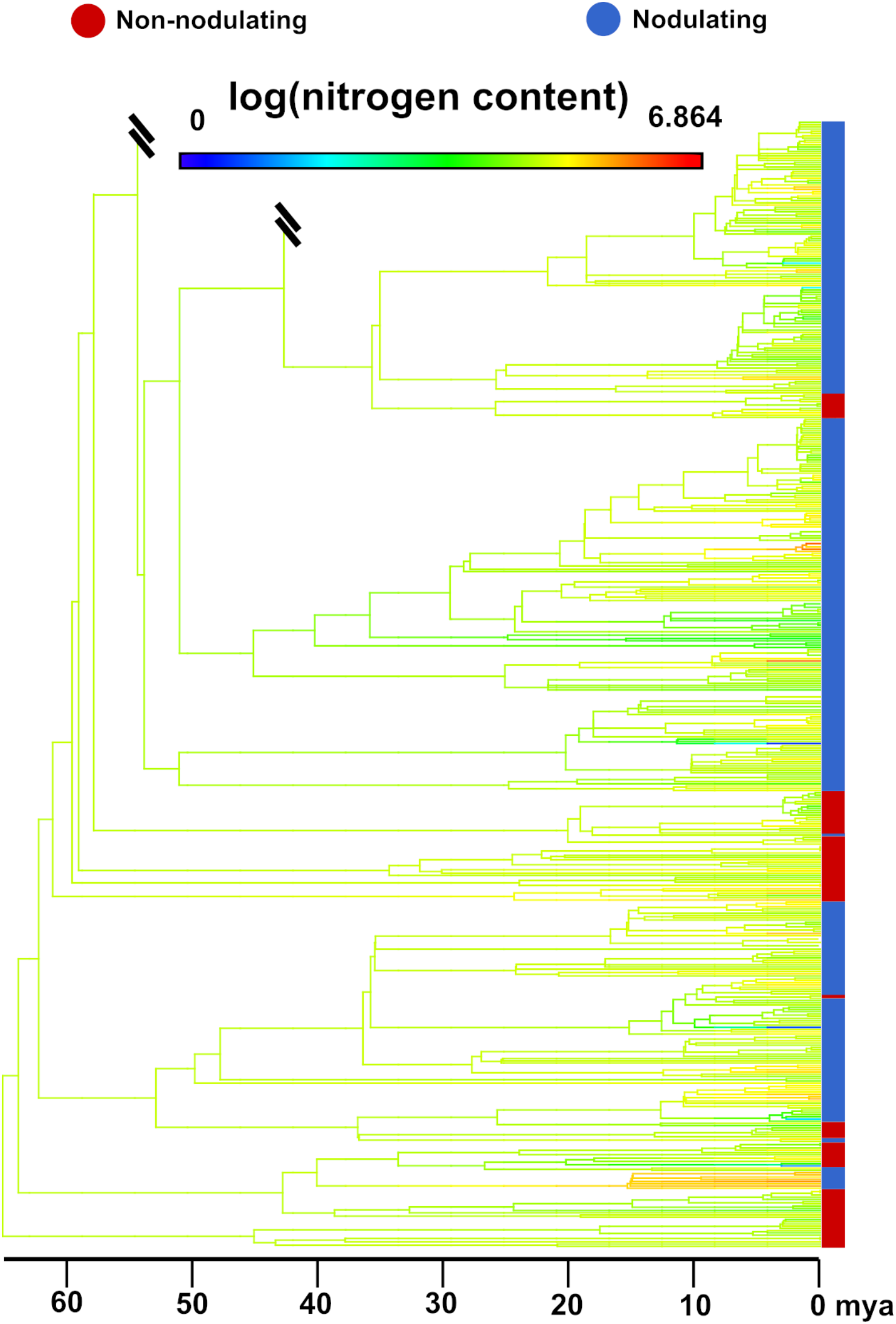
Ancestral reconstruction of nitrogen niche shifts in the early diverging papilionoid legumes, with warmer colors representing richer soil nitrogen. The right-hand side bar indicates species that are either nodulating (blue) or non-nodulating (red). Double hashes represent omitted branches of the total tree (> 6,000 species). Note that while nodulators are overall present in somewhat lower soil nitrogen niche, there are many niche shifts within clades that are constant for nodulation status. The bottom-most blue bar represents the one evident shift in nitrogen niche with concomitant nodulation (genus *Dussia*, a neotropical rainforest clade), but this is into richer nitrogen niche.

### Drivers of RNS

RNS has traditionally been interpreted as a direct adaptation to nitrogen-poor soils since its earliest origins (Sprent 1994b, 2009). In contrast, our work corroborates an alternate view (Crews 1999; Pellegrini et al. 2016; Tamme et al. 2021; Doby et al. 2022) that the actual distribution of nitrogen on Earth (LeBauer and Treseder 2008) is weakly associated with nodulation, and other climatic factors are more closely predictive of distributions. Here, the main environmental associate of nitrogen-fixing symbiosis was aridity, as in previous investigations (Pellegrini et al. 2016; Doby et al. 2022).

Aridity being the best predictor aligns with the disproportionate prevalence and diversity of legumes throughout Africa as compared to less legume-dominated environs elsewhere in the tropics (Yahara et al. 2013; Pellegrini et al. 2016; Sprent 2001: p. 8), contrasting with the lower diversity of the African tropical flora as a whole (Terborgh et al. 2016). Similar results were obtained for temperature seasonality, consistent with arguments that nodulators are particularly adapted to seasonally dry tropical areas (Pellegrini et al. 2016). We also found that nodulators were associated with lower mean annual temperature, consistent with the prevalence of nodulating taxa in temperate environments, particularly actinorhizal species and herbaceous legumes of subfamily Papilionoideae (Tamme et al. 2021; Sprent et al. 2013). We also note that rhizobial and actinorhizal plants repeatedly show distinct patterns of environmental tolerance and differing rates of niche evolution, suggesting separate ecological strategies that pair with their distinct phylogenetic histories (Soltis et al. 1995).

That nitrogen was not the best predictor corroborates what has been called the nitrogen-paradox (Crews, 1999), where RNS species are diverse in tropical environments that commonly do not experience nitrogen limitation. As Sprent et al. (2013) and Crews (1999) point out, distinguishing between nodulators and non-nodulators is necessary, as we have done here, as well as a consideration of facultative nitrogen fixation (Sheffer et al. 2015; Hedin et al. 2009; Barron et al. 2011). The mesic and arid tropics contain almost all the non-nodulating diversity of legumes (Sprent et al. 2013), and facultative or even rare RNS in taxa enabled to do so is prevalent (Barron et al. 2011), whereas the almost entirely nodulated papilionoid legumes dominate outside of tropical zones. By contrast to the legumes, the ecology of actinorhizal plants is not well-understood, with the primary biogeographic difference known being that they specialize in temperate environments (Tamme et al., 2021). Causes for diversity being structured differently in temperate and tropical regions is an area of intense interest (Folk et al. 2019; Segovia et al. 2020; Sun et al. 2020; reviewed in Folk et al. 2020) that has received focused study in nodulators (Sprent et al. 2013; Menge et al. 2019; Sprent et al. 2017; Menge and Crews 2016; Menge et al. 2014; Menge et al. 2017; Tamme et al. 2021). Understanding what environmental pressures cause latitudinal differences in the prevalence of RNS, how those pressures play out over time, and how much key traits, such as symbiotic partners and nodule differences in RNS taxa, determine any generalities remains largely unresolved. Future research should move beyond assessing single traits, and consider multiple traits and their associations, to better evaluate latitudinal gradients in diversity and drivers such as resource pressure, symbiont availability, and life history trade-offs.

### Speciation rates

Investigations of nodulation as a diversification-associated key trait have been infrequent (e.g., Lavin et al. 2005). Traditionally, the diversity of legumes has been connected primarily to habitat and biogeographic shifts (Koenen et al. 2013; Pennington et al. 2004; Pennington et al. 2009). Compared to previous investigations (Koenen et al. 2013; Afkhami et al. 2018), we found that actinorhizal and rhizobial RNS have inconsistent diversification responses. Overall, rhizobial nodulators have increased evolutionary rates, while rates of actinorhizal nodulators are markedly lower when compared to non-nodulators (Fig. 2b). This is consistent with the sporadic distribution and poor species richness of extant actinorhizal nodulators (Pawlowski and Sprent 2008; Tamme et al. 2021; Ardley and Sprent 2021). Rhizobial and actinorhizal nodules are structurally very different from one another, and nodules are likewise diverse within each symbiosis type (Pawlowski and Sprent 2008). Combining this morphological and physiological diversity with broader uncertainties about the underlying homology of nodules and the near-certain lack of homology with other types of nitrogen-fixing symbiosis (Doyle 2011, 2016), our results suggest caution in the interpretation of large-scale studies that do not explicitly confront the diversity of RNS.

One candidate for a physiological process underlying the diversification difference between rhizobial and actinorhizal nodulators is efficiency; rhizobial nodules can achieve far higher rates of nitrogen fixation (Pankievicz et al. 2019). Physiologically, actinorhizal nodules differ in their lesser ability to control oxygenic environments, which are hostile to nitrogenase; the bacterial partner *Frankia* instead has intrinsic methods for oxygen exclusion (Pankievicz et al. 2019). However, further investigation of this topic must consider the life history differences among actinorhizal and rhizobial RNS. For instance, actinorhizal nodulators are almost entirely woody plants with relatively long lifespans, whereas herbaceous habits are most common in rhizobial plants.

Rhizobial RNS in the legumes is solely responsible for the high diversification rates seen in nodulators and indeed the entire nitrogen-fixing clade. But even within this large group, speciation dynamics are not uniform. Although the largest legume subfamily, Papilionoideae, has a long right-hand tail of rapid diversifiers, the probability density in the highest speciation rates is in subfamily Caesalpinioideae and specifically the mimosoid clade (Fig. 2c). Mimosoids are most diverse in the dry tropics (e.g., Grass and Succulent biomes, as defined by Lewis et al. 2005), and some form symbiotic associations with the highly distinct branch of betaproteobacteria (*Burkholderia*; Sprent (2009)), while nearly all rhizobial taxa outside this clade are nodulated by alphaproteobacteria (but see Elliott et al. 2007; Rasolomampianina et al. 2005). Alignment of plants with optimal diazotrophs improves symbiosis (Friesen 2012; Heath and Tiffin 2009), and this alignment is dependent on habitat context (Heath and Tiffin 2007). The greater diversity of symbionts recorded in mimosoid species, including co-infection by distinct diazotroph lineages (Barrett and Parker 2006), may therefore suggest a solution to distinctive habitat pressures (Sprent et al. 2017). The number of species investigated for betaproteobacteria symbionts is limited (Sprent 2009), and distinguishing environment and biogeography from host-symbiont interactions is challenging outside of an experimental approach (Heath and Tiffin 2007). Further investigations at lower phylogenetic levels are warranted to test our initial results that symbiont partner is associated with increased diversification rates.

## Conclusions

Here, we used an explicitly phylogenetic approach to address key hypotheses about the possible environmental drivers of RNS and of its potential role in the diversification of the nitrogen-fixing clade. Our results show (1) a key role for aridity, as opposed to nitrogen limitation itself, as the closest environmental associate of RNS state; (2) that life in arid, poor soils represents a stable evolutionary strategy for nodulators; and (3) early transitions to nodulation followed by later entry into habitats favoring nodulation. We also identified different ecological and evolutionary patterns in rhizobial versus actinorhizal RNS, which have distinct environmental niches and speciation dynamics. In addition, subclades within rhizobial legumes showed differing speciation rates, perhaps related to different bacterial symbiont taxa. Finally, one of our most surprising results was the lack of temporal coincidence between early gain of nodulation and late shifts into environments favorable for the maintenance of nitrogen-fixing symbiotic strategies. These results point to nodulation as an exaptation that facilitated subsequent shifts into arid, poor soils.

## ACKNOWLEDGMENTS

We thank Euan James for guidance on assigning nodulation states and Jean-Michel Ané for discussions on the evolution of nitrogen fixation. We also thank herbarium curators and staff of BRIT, CAS, FLAS, F, HUH, KUN, MO, NY, OS, TEX, US for facilitating the major sequencing initiative that enabled this project.

## AUTHOR CONTRIBUTIONS

CMS, RAF, RPG, PSS, DEG designed the research, CMS, RAF, HRK carried out the research, CMS and RAF conducted data analysis, RAF and CMS wrote the manuscript and all authors read, commented, and approved it before submission.

## DATA AVAILABILITY

Scripts and data to perform the above analyses are posted on GitHub (https://github.com/carol-siniscalchi/NitFixSoil). Nodulation data and backbone constraints are available from Kates et al. (2022).

